# High-resolution Inference of Multiplexed Anti-HIV Gene Editing using Single-Cell Targeted DNA Sequencing

**DOI:** 10.1101/2024.01.24.576921

**Authors:** Mohamed S. Bouzidi, Zain Y. Dossani, Carolina Di Benedetto, Kyle A. Raymond, Shivani Desai, Leonard R. Chavez, Paola Betancur, Satish K. Pillai

## Abstract

Gene therapy-based HIV cure strategies typically aim to excise the HIV provirus directly, or target host dependency factors (HDFs) that support viral persistence. Cure approaches will likely require simultaneous co-targeting of multiple sites within the HIV genome to prevent evolution of resistance, and/or co-targeting of multiple HDFs to fully render host cells refractory to HIV infection. Bulk cell-based methods do not enable inference of co-editing within individual viral or target cell genomes, and do not discriminate between monoallelic and biallelic gene disruption. Here, we describe a targeted single-cell DNA sequencing (scDNA-seq) platform characterizing the near full-length HIV genome and 50 established HDF genes, designed to evaluate anti-HIV gene therapy strategies. We implemented the platform to investigate the capacity of multiplexed CRISPR-Cas9 ribonucleoprotein complexes (Cas9-RNPs) to simultaneously 1) inactivate the HIV provirus, and 2) knockout the *CCR5* and *CXCR4* HDF (entry co-receptor) genes in microglia and primary monocyte-derived macrophages (MDMs). Our scDNA-seq pipeline revealed that antiviral gene editing is rarely observed at multiple loci (or both alleles of a locus) within an individual cell, and editing probabilities across sites are linked. Our results demonstrate that single-cell sequencing is critical to evaluate the true efficacy and therapeutic potential of HIV gene therapy.

## Introduction

Despite the advent of antiretroviral therapy (ART), the human immunodeficiency virus (HIV) was responsible for the deaths of 650,000 people worldwide in 2021, while 38.4 million individuals are currently living with HIV infection.^1^ Moreover, ART needs to be taken on a lifelong basis due to the persistence of HIV latently-infected cells during treatment, which typically reinitiate spreading infection when ART is stopped. ^2,3^ As access to lifelong ART is extremely challenging in resource-limited settings where HIV predominates, there is a desperate need for effective prophylactic and curative strategies. ^4,5^

Several HIV cure strategies are currently in development, including “shock-and-kill”,^6^ “block-and-lock”^7^, and oncology-derived immunomodulatory ^8^ and bone marrow transplantation approaches targeting the latent HIV reservoir. ^9–11^ The discovery of the CRISPR (Clustered Regularly Interspaced Short Palindromic Repeats)-Cas9 gene editing system has raised new hopes for gene therapy-based HIV cure strategies.^12–14^ The Cas9 enzyme interacts with the scaffold sequence of a guide RNA (gRNA) complementary to a target sequence, enabling knockout of a targeted gene in a highly specific manner.^15^ HIV gene therapy-based cure strategies in development typically aim to excise the HIV provirus directly,^16–22^ or target host dependency factors (HDFs) including viral entry receptors that support viral persistence.^23–25^ In recent years, a number of preclinical HIV cure studies have showed promising results using this technology. ^17,19,26–29^ An important caveat, however, arises from the high mutation and recombination rate of HIV-1,^30,31^ which allows the virus to navigate broad expanses of sequence space in short periods of time. Studies of CRISPR-based HIV excision strategies targeting the viral genome at a single site demonstrate rapid evolution of viral resistance to the gene editing modality, mirroring the evolution of antiviral drug resistance in the setting of monotherapy.^32–36^ Therefore, the HIV genome will need to be targeted at multiple sites simultaneously. Drawing from lessons learned during the development of combination ART, a minimum of three distinct genome sites may need to be targeted to prevent evolution of resistance.^16^ Similarly, the neutralization of a single HDF supporting HIV replication is unlikely to completely protect cells from infection, suggesting that multiple HDFs will likely need to be targeted simultaneously to achieve a functional or sterilizing cure for HIV infection.^16^ Even when considering a single HDF locus, genes exhibiting codominance between alleles will likely require biallelic knockout to fully eliminate generation of the encoded protein and, in this manner, block viral infection. Illustration of this concept is provided in nature; CCR5-Δ32 heterozygotes typically produce enough CCR5 protein at the target cell surface to support HIV replication, while Δ32 homozygotes are fully protected from HIV infection (by the highly prevalent CCR5-tropic HIV strains).^37–39^

Based on the need for multitargeting to achieve an HIV cure, molecular diagnostics enabling detection of multiple edits within individual cells will be critical in assessing the true, real-world potency of anti-HIV gene therapy approaches. Currently-implemented technologies, including Sanger sequencing of bulk cell populations and gel-based imaging of excision products, do not enable inference of co-editing within single cells.^40^ Furthermore, standard deep sequencing approaches will not suffice, due to read length limitations that prevent inference of genetic linkage between distal sites.^40^ To address this need, we report here a novel, targeted single-cell DNA sequencing (scDNA-seq) pipeline featuring a hybrid experimental and computational structure to characterize anti-HIV gene editing at viral and host genome sites with extreme resolution. Our platform characterizes the near full-length HIV genome and 50 established HDF genes (identified in a primary cell-based CRISPR screen) at the single-cell level. We implemented the platform to investigate the capacity of multiplexed CRISPR-Cas9 ribonucleoprotein complexes (Cas9-RNPs) to simultaneously 1) inactivate the HIV provirus, and 2) knockout the *CCR5* and *CXCR4* HDF (entry co-receptor) genes in microglia and in primary monocyte-derived macrophages (MDMs), leveraging our previously published work^29^. Our findings demonstrate the utility and critical importance of scDNA-seq analysis to accurately and comprehensively evaluate the efficacy of antiviral gene editing strategies.

## Results

### Generation and validation of a customized panel allowing detection of CRISPR-Cas9 edits in the near-full length HIV provirus and HIV HDFs at the single-cell level

To develop our single-cell sequencing strategy to detect CRISPR-Cas9 editing, we first designed a sequencing primer panel. To evaluate gene editing within both the HIV provirus and HDFs supporting viral replication, we designed a panel targeting tiled amplicons spanning the HIV genome, and 53 HDF genes that were previously identified using a genome-wide CRISPR screen in primary CD4+ T cells (e.g. *CXCR4*, *CCR5* and *CD4*, which also facilitate HIV entry into myeloid cells).^41^ All targeted genes, chromosomal positions, primer pair names, sequences, and genome coordinates are described in the supplementary materials (Supp. Table 1). Thirty-six primer pairs cover the HIV genome from position 30 to position 9516 based on the HIV HXB2 reference genome (Figure 1). This includes the 5’ long terminal repeat (LTR) region, which has been targeted specifically in several HIV gene therapy strategies,^16,17,19,26–28,42^ allowing detection of edits in highly conserved regions such as the Primer Binding Site (PBS) and packaging signals (PSI sequences).

**Figure 1.**
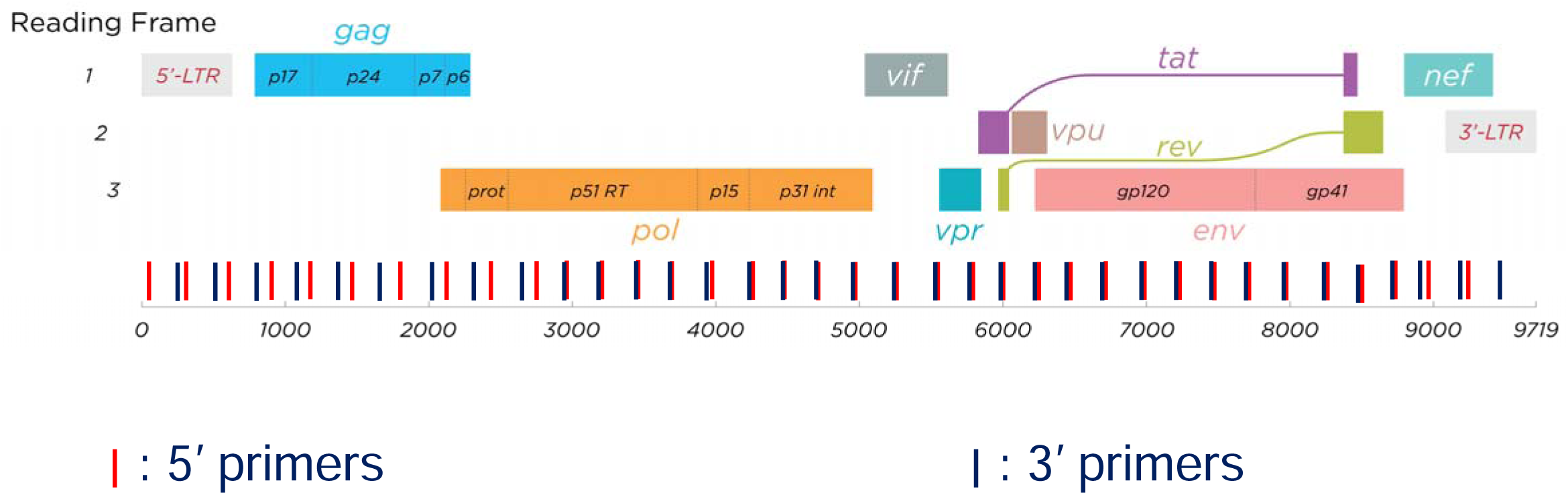
Map of primer binding sites across the HIV genome. Open reading frames across the entire HIV genome are represented. Blue and red bars represent forward and reverse sequencing primers, respectively. A total of 36 primer pairs were employed to achieve near-complete coverage of the HIV genome. Nucleotide coordinates corresponding to each primer pair are provided below the bars.

### Bulk cell and single-cell sequencing analyses provide concordant estimates of antiviral gene editing efficiencies

In order to test our single-cell sequencing approach, we first targeted the HIV 5’LTR along with the host HIV co-receptor genes *CXCR4* and *CCR5* using CRISPR-Cas9 RNP complexes. We first tested our approach in the HC69.5 cell line, a clone derived from the C20 immortalized microglial cell that has been infected with an HIV-derived construct.^43^ The presence of a d2eGFP sequence under the control of the viral promoter permits the detection of viral expression through flow cytometry measurement of GFP signal. Moreover, no latency reversal agent treatment is required to detect the GFP signal as the HC69.5 cell line expresses HIV constitutively.^44^ We nucleofected HC69.5 cells with RNPs targeting the HIV 5’LTR, *CXCR4* and *CCR5* and evaluated editing efficiency using our scDNA-seq platform, Sanger sequencing and Tracking of Indels by Decomposition (TIDE) analysis, and flow cytometry (Figure 2A).

**Figure 2.**
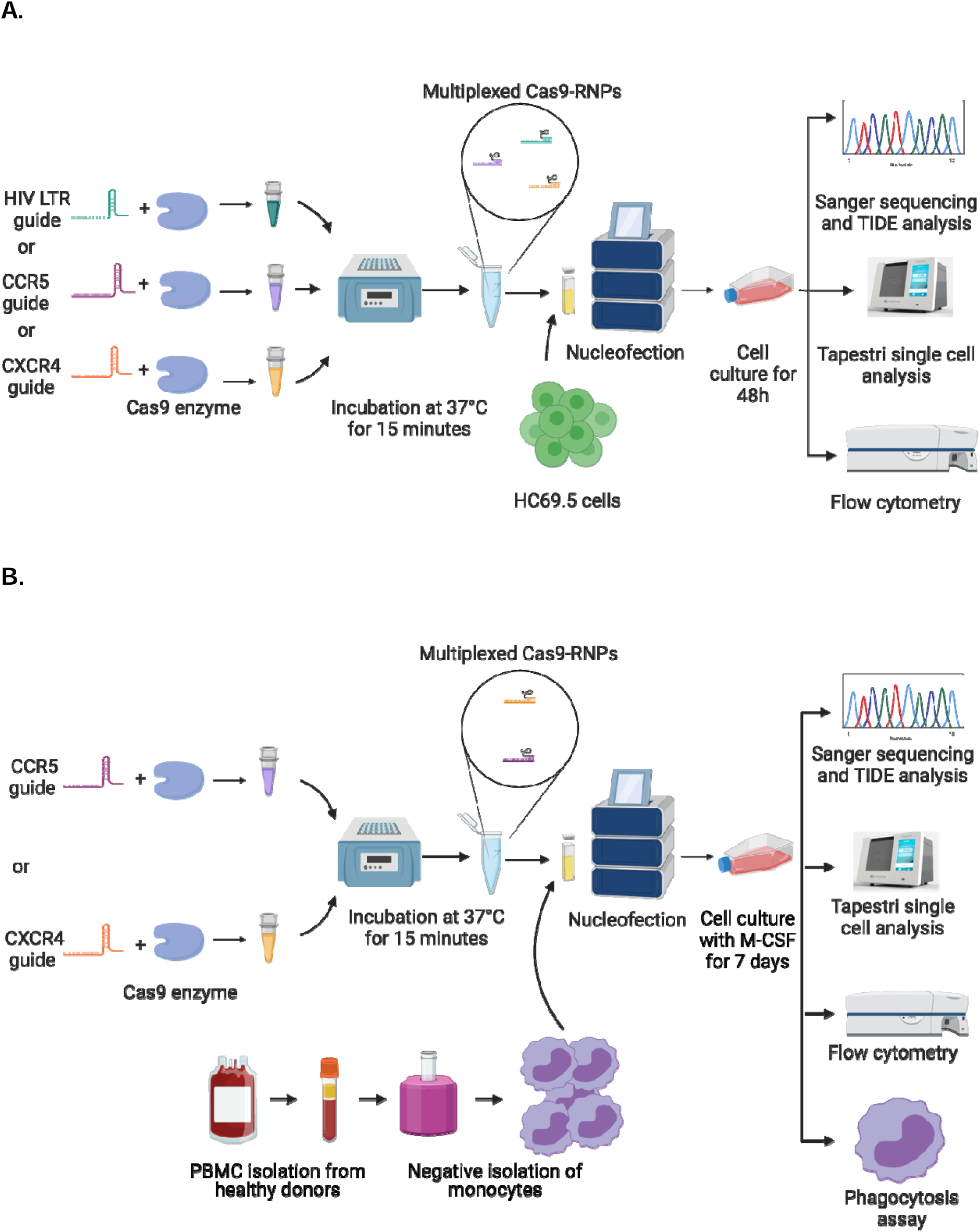
Schematic of anti-HIV gene editing workflow. A) RNP complexes were generated using guides targeting the LTR, CXCR4, or CCR5 loci. The resulting complexes were pooled at equimolar concentrations and delivered to HC69.5 cells utilizing a Lonza 4D-Nucleofector. After 48 hours of culture, cells were harvested for flow cytometry, DNA isolation, PCR and TIDE analysis and Tapestri single cell analysis. B) PBMCs were isolated from whole blood obtained from healthy donors. Monocytes were then isolated using a negative selection antibody cocktail. Simultaneously, CXCR4 and CCR5 targeting guides were generated and complexed with purified Cas9 protein to form CXCR4 and CCR5 targeting Cas9-RNPs. These RNPs were pooled at equimolar concentrations and delivered to freshly isolated monocytes using a Lonza 4-D Nucleofector. Following stimulation with M-CSF for 7 days, MDMs were polarized. Harvested cells underwent DNA extraction for PCR amplification and Sanger sequencing, scDNA-seq library preparation, flow cytometry, and phagocytosis assays.

In regards to coverage across the HIV genome (Supplementary Figure 1A), we observed a gap in coverage between amplicons 6 and amplicon 21. This reflects the fact that the provirus in the HC69.5 clone lacks the corresponding region of the HIV genome (consisting of a part of *gag* and the whole *pol* gene). We independently evaluated and validated the performance of our panel by applying our scDNA-seq pipeline to the J-Lat cell line (Jurkat cells harboring a near-complete, latent HIV proviral genome). The read depths observed across amplicons indicate robust coverage across the HIV provirus (Supplementary Figure 1B). Similar metrics were generated for the host genes indicating robust coverage (Supplementary Figure 1B and C).

We next evaluated gene editing efficiencies at viral and host loci using both Sanger (bulk) and single-cell NGS approaches. Three measurements were generated for each targeted site: 1) TIDE analysis of Sanger bulk sequencing data, 2) Pseudo-bulk analysis of single-cell NGS data using CRISPResso2 software (ignoring single-cell barcodes), and true single-cell analysis using CRISPResso2 (interpreting single-cell barcodes). Overall, the three distinct analytical methods (leveraging two distinct sequencing approaches) provided highly concordant estimates of gene editing efficiencies at all three loci (Figure 3). We observed high editing efficiencies in the HIV LTR and *CXCR4* (mean efficiency of 74.75 and 60.7%, respectively), while editing of *CCR5* as measured by all three methods was less efficient (21.11% mean editing efficiency) (Figure 3A).

**Figure 3.**
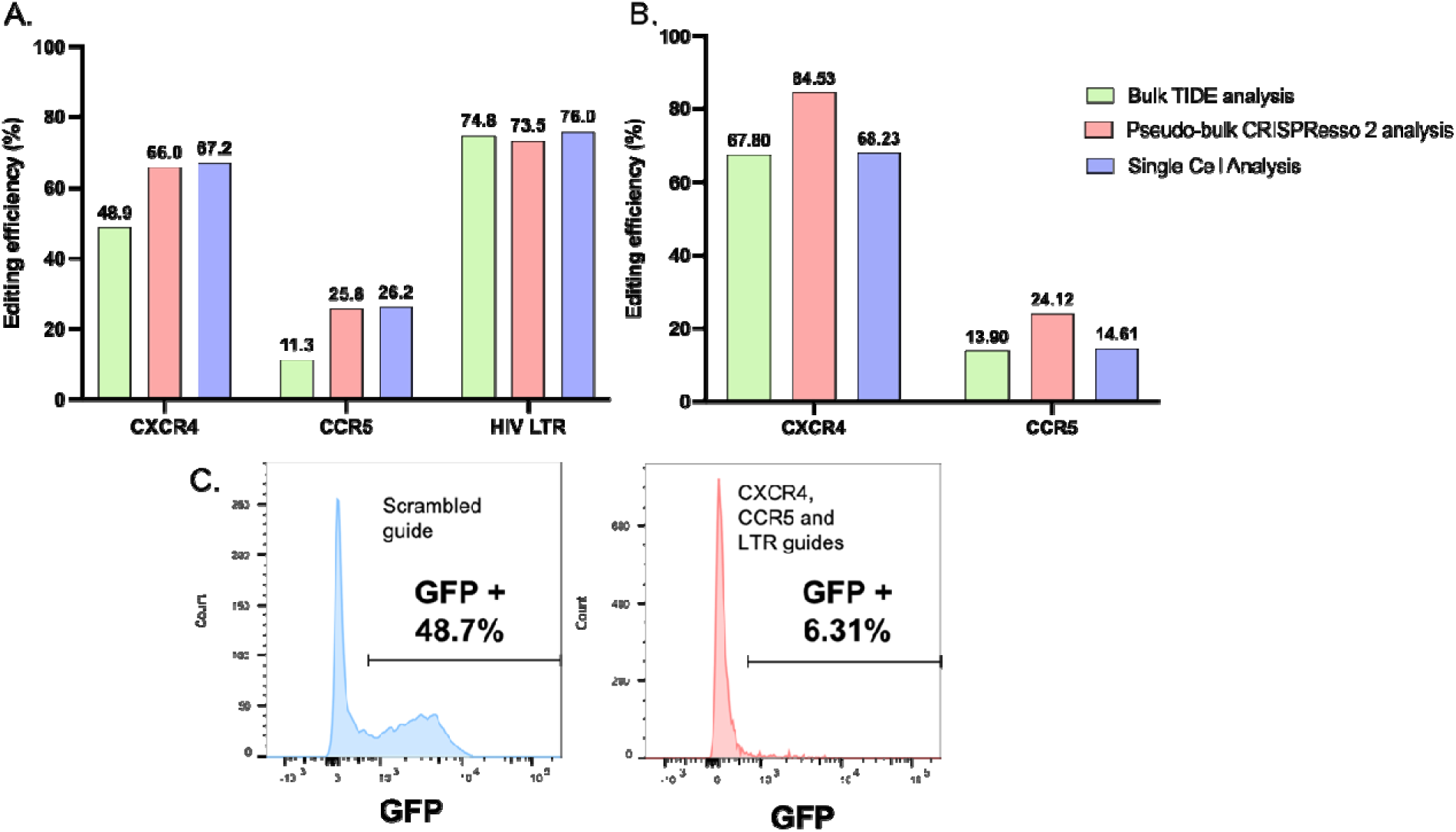
Comparison of inferred gene editing efficiencies between single-cell and bulk Sanger sequencing approaches. Inferred editing efficiency comparison obtained using bulk TIDE analysis (blue), bulk CRISPResso2 (red) and single cell CRISPResso2 (green) methods in HC69.5 (A) or primary MDMs (B). Editing efficiencies are indicated above the bars for each method and each target, expressed as percentages. (C) Flow cytometry on HC69.5 cells nucleofected with scrambled guide RNPs (control) or CXCR4, CCR5 and HIV LTR-targeting RNPs. GFP signal indicates HIV LTR activity.

The observed highly potent editing of the HIV LTR was correlated with a 88% reduction of GFP signal in edited cells as compared to cells nucleofected with control guides (48.7% to 6.31%) (Figure 3C), demonstrating the accuracy of our sequence-based inference of gene editing.

In addition to immortalized microglia, we applied our scDNA-seq pipeline to infer antiviral gene editing in primary monocyte-derived macrophages (MDMs), which are key constituents of the latent HIV reservoir *in vivo* that will likely need to be targeted by HIV cure strategies.^45^ We isolated monocytes from freshly-collected blood, and nucleofected them with *CXCR4-* and *CCR5*-targeting RNPs. After a 7-day M-CSF treatment, MDMs were harvested and characterized by genetic and immunophenotypic analyses (Figure 2B). As it is known that CRISPR editing can impact cell function and metabolism, we first sought to evaluate the impact of editing on macrophage immune function. We incubated edited and unedited cells with fluorescent beads and measured phagocytic uptake by macrophages using flow cytometry (Supplementary Figure 2). We observed a similar level of phagocytosis in edited and unedited cells, demonstrating that a key macrophage immune function was not impacted by nucleofection or Cas9 activity.

Similar to our observations in immortalized microglia, bulk-cell and single-cell analyses provided highly concordant estimates of gene editing efficiencies in primary MDMs. We observed high editing efficiency in *CXCR4* (mean efficiency of 73.52%), while editing of *CCR5* as measured by all three methods was once again less efficient (17.54% mean editing efficiency) (Figure 3B).

### Single-cell sequencing enables highly-precise characterization of CRISPR-induced indel spectra

Precisely characterizing the indel spectra induced by CRISPR gene editing is critical to evaluating therapeutic efficacy and safety. It is now known that the CRISPR-induced indel spectrum is correlated with editing efficiency at a specific site; the less diverse the indel spectrum, the higher the editing efficiency.^46^ We therefore sought to determine if our scDNA-seq pipeline enabled higher-resolution characterization of indel spectra as compared to standard Sanger sequencing and TIDE analysis. Our analyses unequivocally demonstrate that scDNA-seq identifies CRISPR-induced indel spectra with significantly enhanced sensitivity (p=0.025, paired t test), typically revealing a broad range of indels and minority variants (Figure 4). In contrast, Sanger sequencing only enables detection of dominant, majority indel variants (Figure 4).

**Figure 4.**
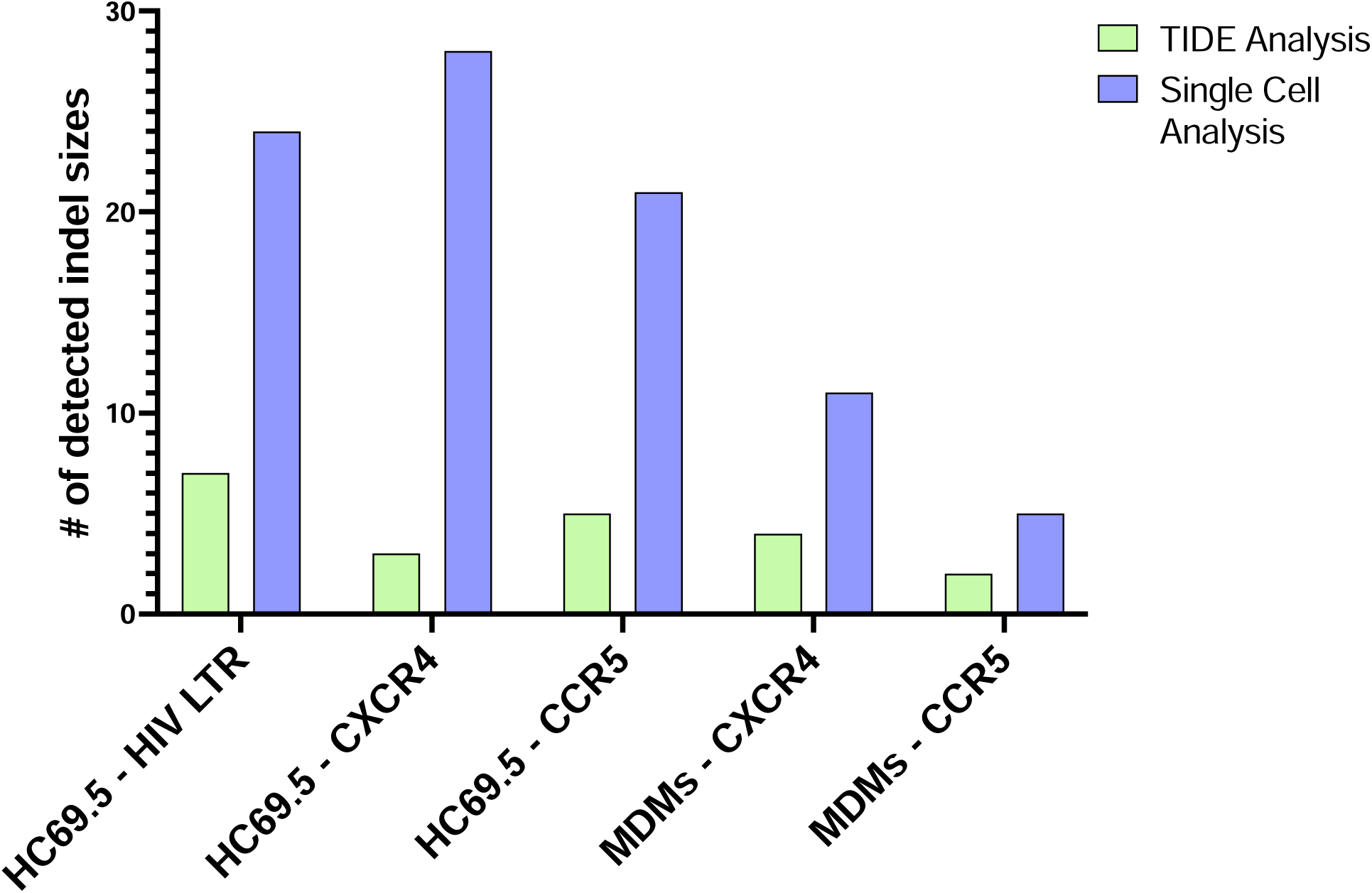
Comparison of inferred indel spectra between single-cell and bulk Sanger sequencing approaches. The sum of the number of peaks inferred using TIDE (green) or single-cell analysis (blue) were plotted for each target in each cell type. The target names and cell types are labeled along the x-axis, with the number of detected indel sizes indicated on the y-axis. Statistical significance was determined using a paired t-test (*p<0.05)

### Single-cell sequencing enables inference of antiviral gene editing at multiple loci within individual cells

HIV enters target cells such as T cells and myeloid cells using the CD4 receptor and the CXCR4 or CCR5 co-receptors. It has been described that a switch occurs during the course of HIV infection, in favor of one or the other co-receptor.^47^ It is therefore necessary to knock-out both co-receptors to avoid a switch, preventing viral rebound. Leveraging our single cell sequencing approach, we sought to evaluate co-editing efficiency at a single-cell level in HC69.5 and primary MDMs. We processed our single-cell sequencing data through CRISPRessoV2 software (as described in the Materials and Methods), and summarized the data in Figure 5.

**Figure 5.**
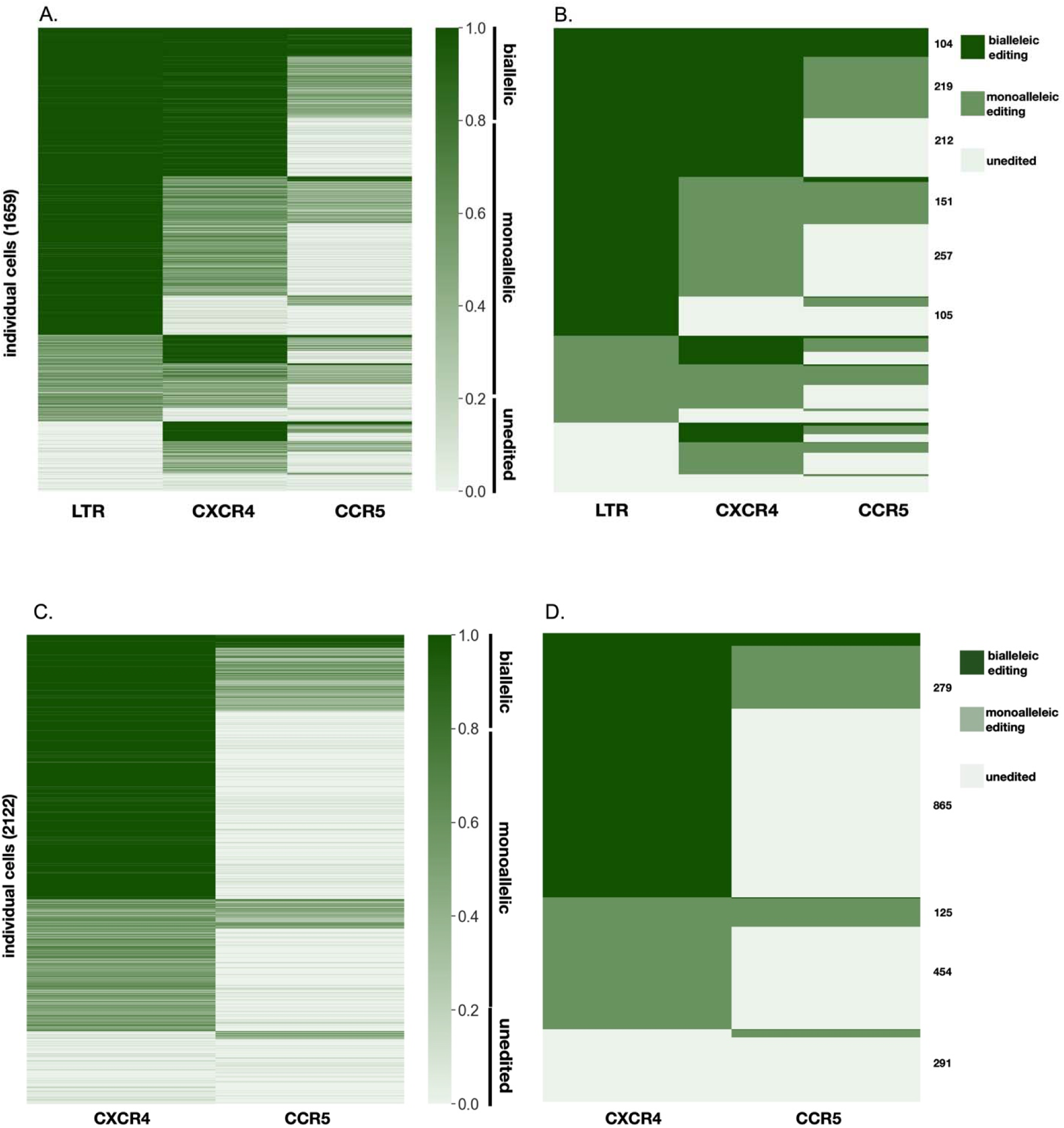
Heatmap visualization of multi-site gene editing within individual cells. Each row in the heatmap represents a single cell, with the shade of green indicating the editing frequency for each target. (A) Detailed view of the editing per cell and per loci analyzed in HC69.5 cells (editing efficiency at each site is represented as a continuous variable). (B) Discretized editing heatmaps for HC69.5 cells. Editing values above 0.8 indicate bi-allelic editing, between 0.2 and 0.8 indicate mono-allelic editing, and below 0.2 indicate no editing. Biallelic editing is represented in dark green, monoallelic editing in light green and unedited in pale green. The number of cells in each group is listed (when counts exceed 100). (C) Detailed view of the editing per cell and per loci analyzed in MDMs. (D) Discretized editing heatmaps for MDMs.

Each line represents one cell, characterized by its editing frequency at each targeted site. The shade of green indicates the editing frequency (dark green for high editing frequency to white for low editing frequency) (Figure 5A&C) Full co-editing (editing of all targets) was observed in 37.4% of HC69.5 cells (Figure 5B) and in 22.38% of primary macrophages (Figure 5D). The single-cell resolution of our analysis demonstrates the extreme heterogeneity of antiviral gene editing patterns across cells (Figure 5), informing therapeutic efficacy while demonstrating that cell-specific factors likely modulate the intake and activity of gene editing machinery.

### Single-cell sequencing enables allele-specific evaluation of antiviral gene editing performance

This single-cell approach also allows us to survey editing in each of the two alleles of each targeted host gene by using specific editing efficiency thresholds.^48^ This is of particular importance as editing on one chromosome may not be enough to fully shut down the expression of a HDF and prevent HIV infection. For this purpose, we categorized allele-specific editing based on observed editing efficiencies (defined as the frequency of NGS reads harboring indels); genes with an editing efficiency above 0.8 were considered to be edited at both alleles, genes with an editing efficiency value between 0.2 and 0.8 were considered to be edited at only one allele, and genes with an editing efficiency below 0.2 were categorized as non-edited (Figure 5A&C). We observed that 43% of the HC69.5 cells were *CXCR4^mut^*/*CXCR4^mut^*and 42% *CXCR4^wt^*/*CXC4^mut^*. For CCR5, we observed that 10% of the cells *CCR5^mut^*/*CCR5^mut^* and 36% of the cells were *CCR5^wt^*/*CCR5^mut^* (Figure 5B). In primary macrophages, approximately 57% *CXCR4^mut^*/*CXCR4^mut^*and 28% of the cells were *CXCR4^wt^*/*CXCR4^mut^*. For *CCR5*, we observed that 4% *CCR5^mut^*/*CCR5^mut^*and 21% of cells were *CCR5^wt^*/*CCR5^mut^* (Figure 5D).

### Gene editing events exhibit interdependence

Finally, we leveraged the power of our single-cell inference approach to determine whether antiviral gene editing events occur independently of one another. Briefly, we built contingency tables for each gene pair across both tested cell types (HIV-LTR, *CXCR4* and *CCR5* for HC69.5 and *CXCR4* and *CCR5* for MDMs) and determined if the co-occurrence of editing at both considered sites was observed more or less than expected by random chance. The expected values were determined by multiplying single cell editing efficiencies as measured in Fig. 4 from each considered pair (assuming that there are independent probabilities of editing at each targeted site). We observed that in both HC69.5 and MDMs, mono-editing (editing at a single targeted locus) was lower compared to the expected value, while co-editing and co editing frequency was higher than expected (Figure 6A, Fisher’s exact t-test p-values ranging from 0.003 to <0.0001). Similarly, we sought to determine if editing at two alleles could be explained by independent probabilities. Again, we obsesrved higher than expected frequencies of co-editing, suggesting that antiviral gene editing events are not independent (Figure 6B&C) , p-value ≤ 0.001).

**Figure 6.**
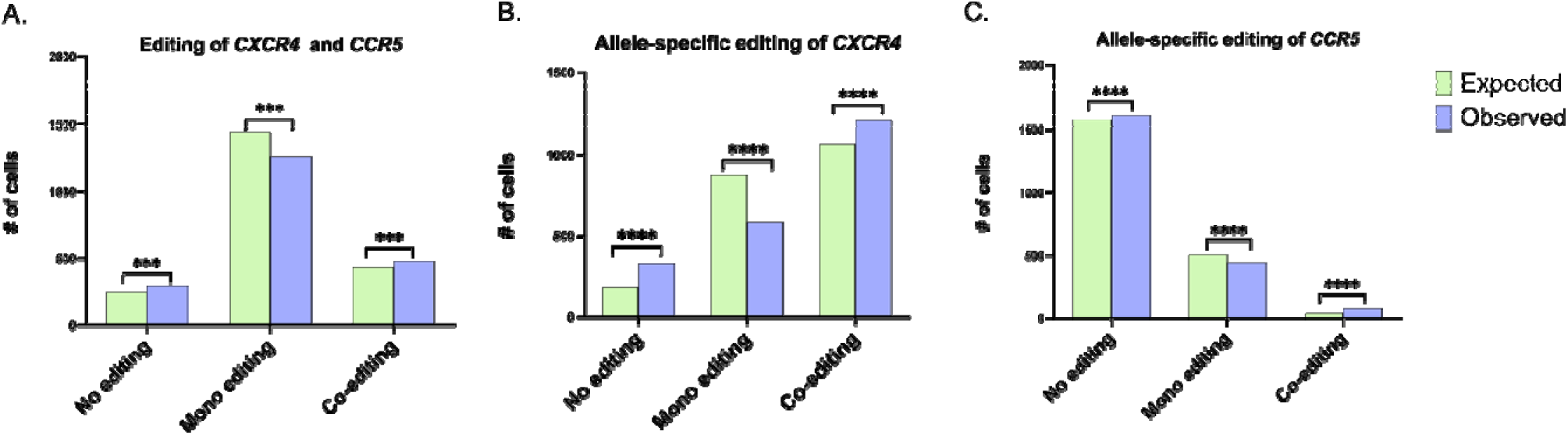
Interdependency of gene editing events. Expected values were determined as the coefficient between the single-cell editing efficiency frequency for one target (as measured in Fig. 4) to the single-cell editing efficiency frequency of the other target (assuming that the probabilities of editing at each targeted site are independent). Expected values of cells harboring no editing, mono editing, or co-editing between CXCR4 and CCR5 loci (A) or between CXCR4 and CCR5 alleles (B & C) have been calculated and plotted (green) and compared to observed values (blue). Statistical significance was determined using Chi-square tests, comparing observed values to expected values (***P < 0.001, ****P < 0.0001).

## Discussion

Anti-HIV gene therapy will likely require editing of multiple host genes and/or sites in the proviral genome. According to mathematical models, more than 99% of total provirus in people living with HIV needs to be inactivated to achieve a functional cure.^16^ Provirus inactivation requires editing on several sites due to viral genetic diversity and plasticity, which may result in the generation of escape mutants considering the exceptional adaptability of HIV.^35^ Moreover, it is unlikely that 99% of infected cells will be targeted by the CRISPR-Cas9 system: some macrophages for example can infiltrate deep into tissues,^49^ where delivery of the CRISPR payload can be very challenging.^50^ One way to overcome this issue is to target HDFs such as HIV co-receptors *CXCR4* and *CCR5* to prevent viral spread from cells harboring unedited provirus. ^51–54^ In this case, mathematical models predict that inactivating provirus in 60% of the infected cells and reducing the pool of susceptible cells by 68% is enough to achieve a functional cure.^16^

So far, no method has been proposed to rigorously assess the frequencies of such events *in vitro* or *in vivo*. Here, we demonstrate that single-cell encapsulation and targeted next generation sequencing, in combination with a specific bioinformatic pipeline, is a reliable and powerful method to characterize antiviral gene editing in both cell lines and primary cells. The editing efficiencies measured with our single-cell method are close but not identical to the results obtained with Sanger sequencing and TIDE analysis. The difference observed between the two methods can likely be explained by the reduced sensitivity of Sanger sequencing in comparison to NGS approaches,^55^ resulting in minority genetic variants being excluded from the final results. Interestingly, the editing efficiency in cells targeted with only one guide at a time is similar to editing obtained in our multiplexed approach, likely due to saturation effects (data not shown).

Highly precise CRISPR editing at a targeted site is key to avoid lethal editing or production of aberrant cells. It has been shown that indel spectra directly correlate with editing precision and efficiency; the less diverse the indels, the more precise and potent the editing.^46^ However, Sanger sequencing and TIDE analysis do not capture indel spectra with sufficient sensitivity, as revealed by our analyses. We demonstrated that indel spectra characterized with our single-cell approach were significantly more detailed and reliable than spectra obtained through TIDE, revealing several indel variants that were unseen in population sequencing data. This result is of particular importance when evaluating a therapeutic approach to predict gene editing efficiency and overall editing precision.

Single-cell sequencing is the only method that enables inference of co-editing events within the same targeted cell. Our results demonstrate that the frequency of co-editing events cannot simply be predicted by considering the product of editing efficiencies across individual loci due to linkage effects. Single cell sequencing also allows us to determine if gene editing is bi-allelic or mono-allelic; this is critical as neutralization of both alleles is often required to completely abrogate target protein expression. In this study, we were able to achieve 42% mono-allelic editing and 43% bi allelic-editing on the *CXCR4* gene in HC69.5 cells. For the *CCR5* gene, 36% of the cells showed mono-allelic editing and 10% bi-allelic editing. The results for *CCR5* reveal a considerable bias towards mono-allelic editing for this locus. Similar results were observed for primary MDMs, where bi-allelic editing of *CXCR4* was observed in 57% versus 28% for mono-allelic editing. For *CCR5*, 21% of the cells were edited in one allele and 3.5% of the cells in both alleles. Interestingly, flow cytometry data for the latter case showed a decrease of 11.3% in the frequency of CCR5-expressing cells compared to the control, and a 13% decrease in mean fluorescence intensity (MFI) (data not shown), without any impact on cell function (as represented by phagocytic activity). This demonstrates that the relationship between gene editing and protein expression is complex, and in some cases, the intact allele may compensate for the knockout allele resulting in a lower-than-expected impact on protein expression. These results highlight the need for a comprehensive method to evaluate gene editing activity such as the single cell approach used here.

The fact that our observed gene editing frequencies across loci and alleles on a cellular level were not independent suggests that some cells may be more permissive than others to editing. The permissiveness of a cell to Cas9-RNPs could be due to several factors: the efficiency of nuclear transport machinery, epigenetic factors such as chromatin state or methylation, or DNA repair machinery.

In conclusion, our study demonstrates that our single-cell pipeline is a powerful approach to evaluate the potency of multi-targeting antiviral gene editing strategies. It yields critical insights into co-editing events across targeted loci and alleles, of key relevance to anti-HIV gene therapy. Our method and panel design can be further expanded to survey a broad range of bioinformatically-predicted off-target sites, to determine the specificity and safety of gene therapy approaches. In addition, our panel can be expanded to reliably characterize non-subtype B proviruses, which will be an essential developmental step when evaluating therapeutic efficacy in diverse global settings. In combination with rational guide design and continuing innovation in the engineering of gene editing machinery, our single-cell sequencing approach provides another key tool to facilitate the clinical development of cure-focused HIV gene therapy approaches.

## Materials and Methods

### Primary Human Monocyte Isolation

Primary monocytes were isolated from blood obtained from healthy donors (Vitalant, San Francisco, CA). A total of 15 mL of blood from TRIMA residual was diluted at a 1:1 ratio with a 3mM EDTA PBS 2% FBS buffer. The resulting mixture was layered onto a Ficoll-Paque density gradient medium and centrifuged at 1200 g for 30 minutes at room temperature. The peripheral blood mononuclear cell (PBMC) layer was carefully collected and subsequently resuspended in 500 µL of ACK lysis buffer (Gibco), followed by a 3-minute incubation at room temperature. To this, 40 mL of 3mM EDTA PBS 2% FBS buffer was added, and the cells were centrifuged for 2 minutes at 1200g. The PBMCs were then subjected to two washes with 50 mL of 3mM EDTA in PBS with 2% FBS, each followed by a 2-minute centrifugation at 1200g. After cell counting, 5 × 10^7 PBMCs were resuspended in 8.5 mL of 3mM EDTA in PBS with 2% FBS. Monocytes were subsequently isolated using the EasySep™ Human Monocyte Isolation Kit (StemCell Technologies) in accordance with the manufacturer’s protocol.

### Cell culture and Macrophage differentiation

The HC69.5 cells were cultured in Dulbecco’s Modified Eagle Medium (DMEM) (ThermoFisher Scientific, Cat. # 11965-118) supplemented with 10% fetal bovine serum (FBS) (Corning, Inc., Cat. #35-010-CV) and 1% Penicillin/Streptomycin (Fisher Scientific, Cat. #11548876) at 37°C with 5% CO2.

Freshly isolated monocytes were seeded in 12-well plates with 800 µL of differentiation medium comprising ImmunoCult™-SF Macrophage Medium (StemCell Technologies) supplemented with 50 µg/mL human M-CSF (PeproTech) for macrophage differentiation. After 7 days, fully differentiated cells were harvested using StemPro Accutase (Gibco) for downstream analysis.

### RNP preparation and cell nucleofection

CRISPR RNA (crRNA) were designed *in silico* using the IDT Design Tool (Integrated DNA Technologies) based on their on-target and off-target score (highest score and lowest score, respectively). The sequences are for *CXCR4* : 5’-GAAGCGTGATGACAAAGAGG-3’, CCR5: 5’-CATCATCTATGCCTTTGTCG-3’ and HIV LTR: 5’-CTACAAGGGACTTTCCGCTG-3’. A mixture of 160 µM tracrRNA (Integrated DNA Technologies) and 160 µM crRNA (Integrated DNA Technologies) was incubated at 37°C for 30 minutes to generate guide RNA (gRNA) at a concentration of 80 µM. Subsequently, 4 µL of gRNA was combined with 4 µL of purified SpCas9-NLS protein (QB3 core, Berkeley) at 40 µM and incubated at 37°C for 15 minutes to form Cas9-Ribonucleoproteins (Cas9-RNPs). For nucleofection, 8 µL of Cas9-RNPs were nucleofected per 1 × 10^6 HC69.5 cells or 1 × 10^6 monocytes, following the manufacturer’s recommendations for a Lonza Nucleofector. The DN-100 program with SF Buffer was employed for HC69.5 cells, while the DS-150 program with P3 buffer was utilized for monocytes.

### DNA Extraction, Sanger Sequencing and TIDE analysis

Cells were harvested, and genomic DNA was extracted from approximately 0.5 × 10^6 cells using Qiagen Blood and Tissue Kits following the manufacturer’s recommendations. The CRISPR-targeted sequences were subsequently amplified using Phusion® High-Fidelity DNA Polymerase (NEB). The primers used for the PCR were as follows: for *CXCR4*, 5’-AGAGGAGTTAGCCAAGATGTGACTTTGAAACC-3’ and 5’-GGACAGGATGACAATACCAGGCAGGATAAGGCC-3’; for *CCR5*, 5’-CTCACTATGCTGCCGCCCAGTG-3’ and 5’-TCACCAGCCCACTTGAGTCCGT-3’; and for HIV LTR, 5’-GCAAGTAGAAGAGGCCAATGAAGG-3’ and 5’-TGCTAGAGATTTTCCACACTGACTA-3’. The PCR products were purified using NucleoSpin Gel and PCR Clean-Up kits (Macherey-Nagel) and subsequently sent for Sanger sequencing (ElimBio, California). AB1 sequencing files were then analyzed using TIDE software ^56^.

### Flow Cytometry

For GFP measurement, HC69.5 cells were harvested and washed three times in PBS (Gibco). Cells were resuspended at a concentration of 1 × 10^6 cells per mL and analyzed using an LSRII flow cytometer. For CCR5 flow cytometry, approximately 1 × 10^6 cells were washed three times with PBS and resuspended in 100 µl of cell staining buffer. Cells were incubated with 5 µl of Human TruStain FcX™ to block Fc receptors for 15 minutes, followed by APC anti-human CD195 (CCR5) (clone J418F1, BioLegend) or 5 µl of the corresponding isotype control staining for 30 minutes. Afterward, cells were washed three times with cell staining buffer and resuspended in 500 µl of staining buffer. Samples were run on a BD LSR II flow cytometer (BD Biosciences).

For the phagocytosis assay, macrophages were washed and resuspended in serum-free IMDM at a concentration of 1 × 10^6 cells/mL. A mixture of 100 µl of macrophages and 100 µl of pHrodo™ Green E. coli BioParticles (Invitrogen, P35366) at 1 mg/mL was incubated for 45 minutes at 37°C. Phagocytosis was stopped by adding 900 µl of cold FACS buffer (PBS+2% PBS). Cells were centrifuged at 500g for 5 minutes and washed with 1 mL of FACS buffer. Subsequently, cells were resuspended in 100 µl of FACS Buffer and incubated with 5 µl of APC anti-human CD14 for 30 minutes. After three washes with FACS Buffer, cells were resuspended in 110 µl of FACS Buffer supplemented with 1 µM Propidium Iodide and analyzed on a FACS Calibur Flow Cytometer (BD Biosciences).

Flow cytometry data were analyzed, and plots were generated using FlowJo (v10.9.0).

### Single-cell library preparation and sequencing

Libraries were prepared following the manufacturer’s recommendations for custom panels (Mission Bio). Briefly, 300,000 HC69.5 or MDMs were encapsulated and barcoded using the Mission Bio Tapestri single-cell sequencing platform. Specific sequences, including HIV LTR, *CXCR4* and *CCR5*, were amplified by a multiplexed targeted PCR using primers described in Supplementary Table 1. The conditions were as follows: incubation at 95°C for 10 minutes, followed by 20 cycles consisting of 30 seconds at 95°C, 10 seconds at 72°C, 6 minutes at 61°C, and 30 seconds at 72°C. Samples were then incubated at 72°C for 2 minutes and kept at 4°C until the next step. All samples exhibited DNA quantities within the expected yield range after PCR clean-up steps. Following library PCR and clean-up, samples were pooled using the Tapestri Sample Quantification Tool, and libraries were sequenced on an Illumina MiSeq using MiSeq Reagent Kit v2 (300-cycles) and a 2x150bp reaction, following the manufacturer’s recommendations.

### Bioinformatic analysis

Sequences were analyzed using a custom pipeline to determine genome editing rates for both the human and HIV genomes at each targeted amplicon in each cell. Briefly, the two 8bp cell barcodes were read from individual sequencing reads and compared to the list of 768 possible barcodes (v1 chemistry) with a custom Python script. Barcodes that were an exact match passed the filter, and those with a Levenshtein distance of 1 away from a unique barcode in the allow list were corrected to that barcode and also passed the filter. Reads that did not have both 8bp barcodes passing the filter were discarded (overall barcode fail rate was 5.3-6.5%). All reads with the same 16bp barcode were collected into individual files. Only barcodes with a minimum of 100 reads were processed using CRISPResso2 in batch mode.^57^ The average number of aligned reads per cell for each amplicon was then plotted. The percentage of reads modified, and the modification state and frequency of *CXCR4*, *CCR5*, and HIV LTR for each cell and each amplicon were analyzed using Python (v3.x) and R Studio (v2023.0.2+561). Briefly, each mutation (indel) was characterized by its size. Cells with fewer than 5 reads for each CRISPR-targeted amplicon were filtered out, and the overall frequency for each indel size and total editing efficiency were plotted.

For the heterozygosity analysis, amplicons corresponding to targeted sites in CXCR4 and CCR5 genes and HIV-LTR were labeled with their mutational state (Wild Type or Mutated), and the frequency of each event was recorded. For each amplicon, each cell was classified into one of three classes: WT/WT, WT/Mut, or Mut/Mut based on the overall percentage of reads with indels in a given cell. Cells with < 20% modified reads were considered WT/WT. Cells with ≥ 20% and < 80% modified reads were considered heterozygous WT/Mut. Cells with ≥ 80% mutated reads were classified as homozygous Mut/Mut. A heatmap has been plotted summarizing these results where each line represents one cell. Custom source code used for the analysis can be found at this address: https://github.com/Pillai-Lab/sc-allelic_antiviral_gene_editing.

### Statistical analysis

The results presented were analyzed with GraphPad Prism v10 (GraphPad Software, SanDiego, CA, USA). Information on specific tests that were used are described in the Results section.

## Acknowledgments

The authors would like to acknowledge Drs. Li Du, Prerna Dabral, and Hannah S. Schwarzer-Sperber at Vitalant Research Institute, and Drs. Robert Durruthy-Durruthy and Shu Wang at Mission Bio for guidance and support in data analysis.

This research was supported by the NIH-funded UCSF-Bay Area Center for AIDS Research (P30 AI027763) specific mechanism: Gilead funded HIV Mentored Scientist award at the amfAR Institute for HIV Cure Research (Vitalant Subaward 12194sc).

## Author Contributions

M.S.B., L.R.C. and S.K.P. initiated the project and designed the experiments. M.S.B., K.A.R. and S.D. performed the bulk sequencing and M.S.B. analyzed the data. M.S.B., K.A.R. L.R.C. performed the single cell sequencing. Z.Y.D. and M.S.B. analyzed the scDNA-seq data. C.D.B. performed the phagocytosis assay. M.S.B. and S.K.P. prepared the manuscript and M.S.B., P.B. and S.K.P. jointly supervised the work.

## Figures and Tables

**Supplementary Figure 1.**
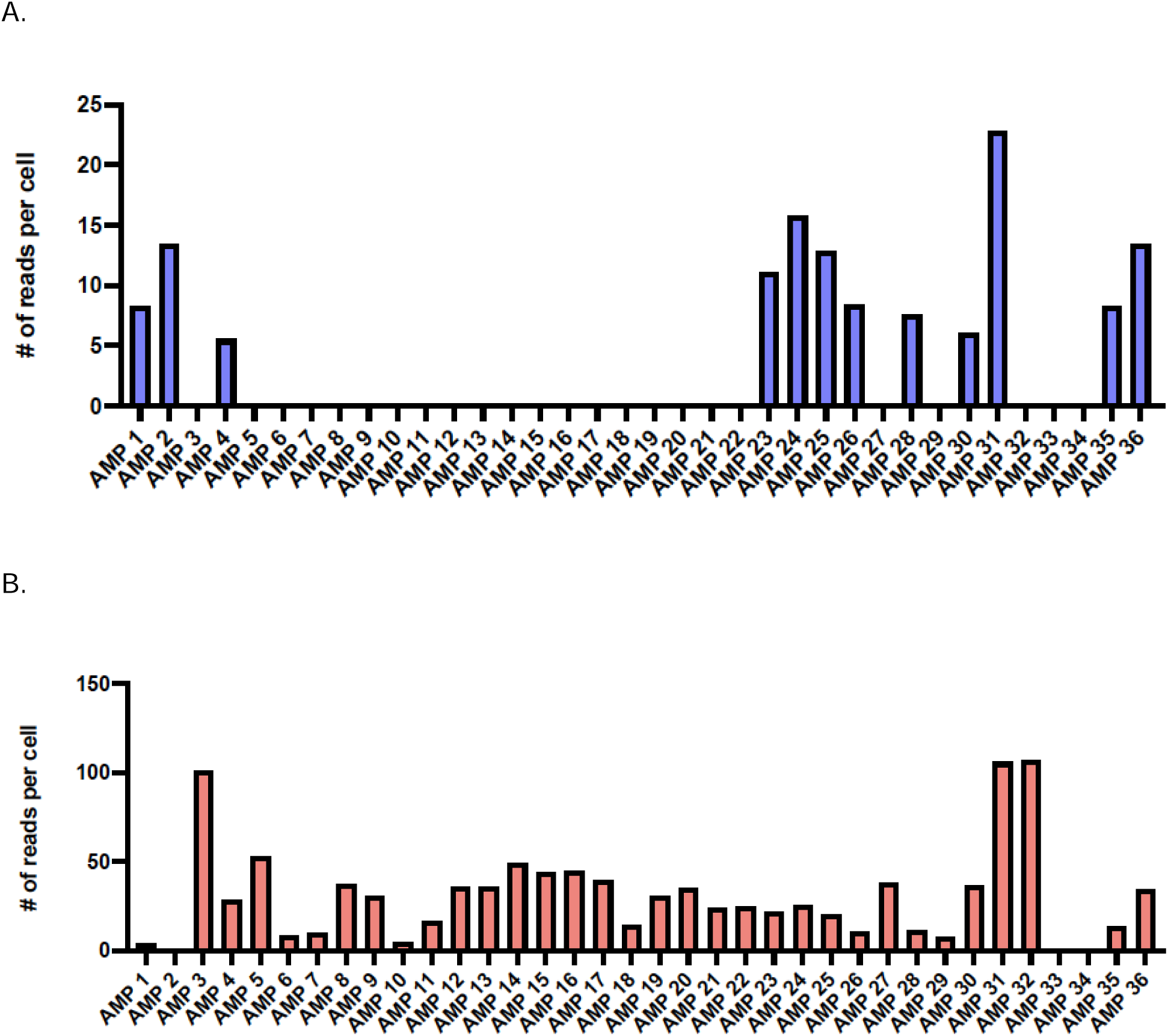
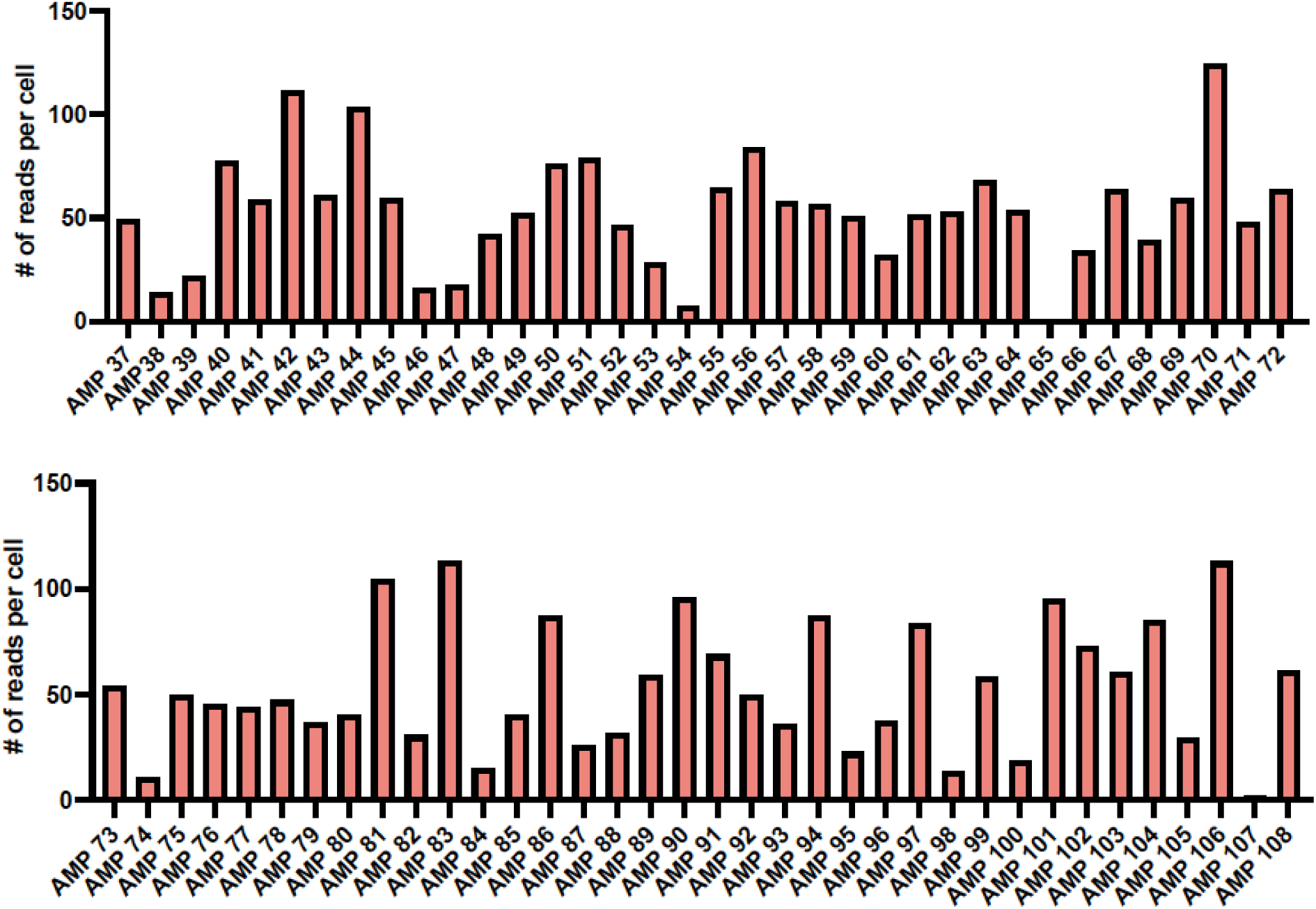
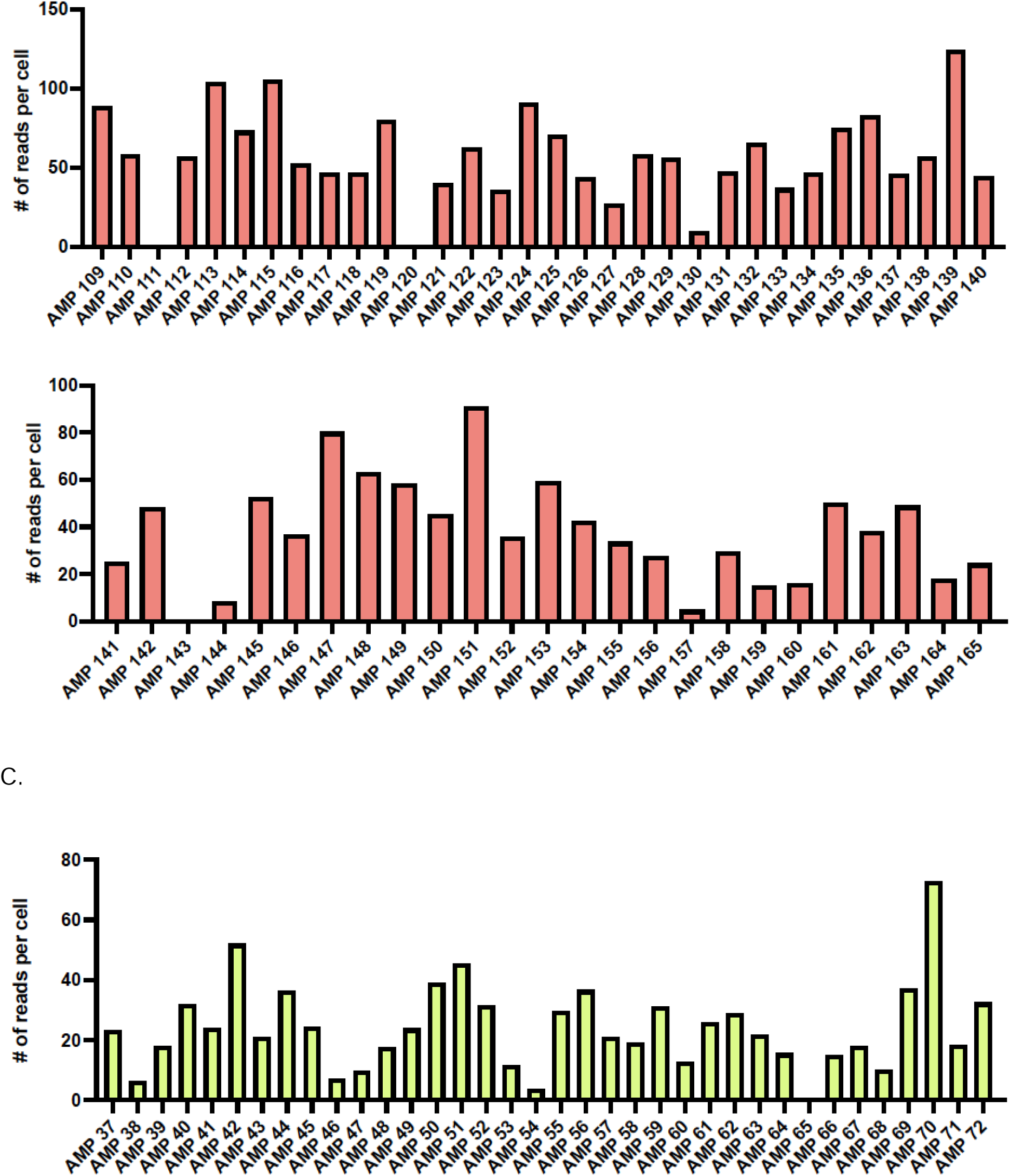
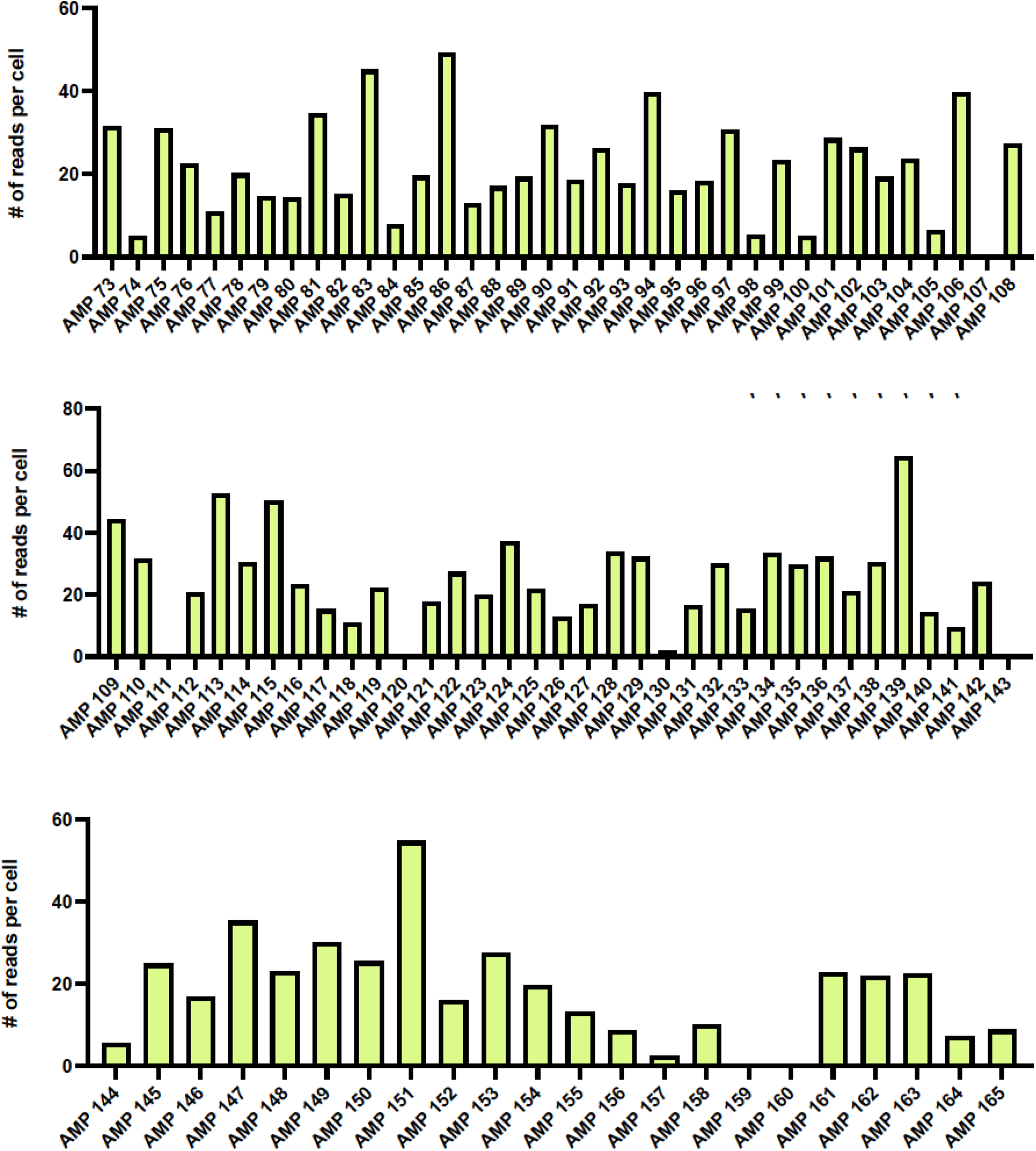
Amplicon coverage across host and viral genome targets. (A) Read coverage for each amplicon in the HC69.5 cell line obtained through the CRISPResso2 pipeline are plotted. The amplicon names are labeled along the x-axis, with the average reads per cell plotted on the y-axis. A gap in reads observed between AMP6 and AMP21 corresponds to the deletion identified in the HC69.5 HIV provirus. (B) Coverage across the HIV genome and host genes in the J-Lat 11.1 cell line. (C) Host gene coverage in HC69.5 cells. The amplicon name is indicated on the x-axis and the average reads per cell is indicated in the y-axis.

**Supplementary Figure 2.**
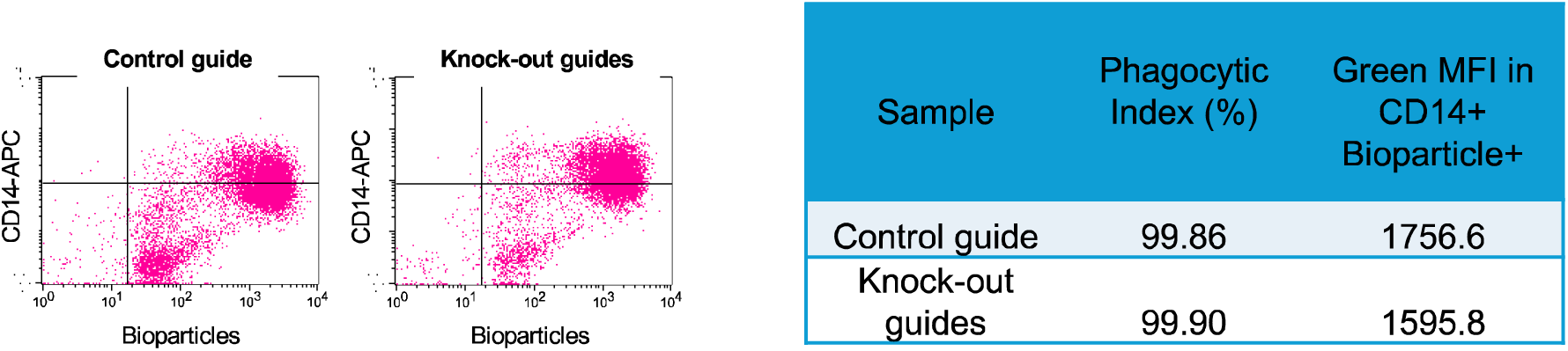
Gene editing workflow does not impede the functional activity of primary MDMs. Cells were incubated for 45 minutes with fluorescent bioparticles to measure phagocytosis and analyzed by flow cytometry. The gating strategy is indicated on the left and the frequency of CD14+ Bioparticles+ cells compared to the total of CD14+ cells and the mean fluorescent intensity (MFI) for control cells and knocked out cells is indicated in the table on the right.

**Supplementary Figure 3.**
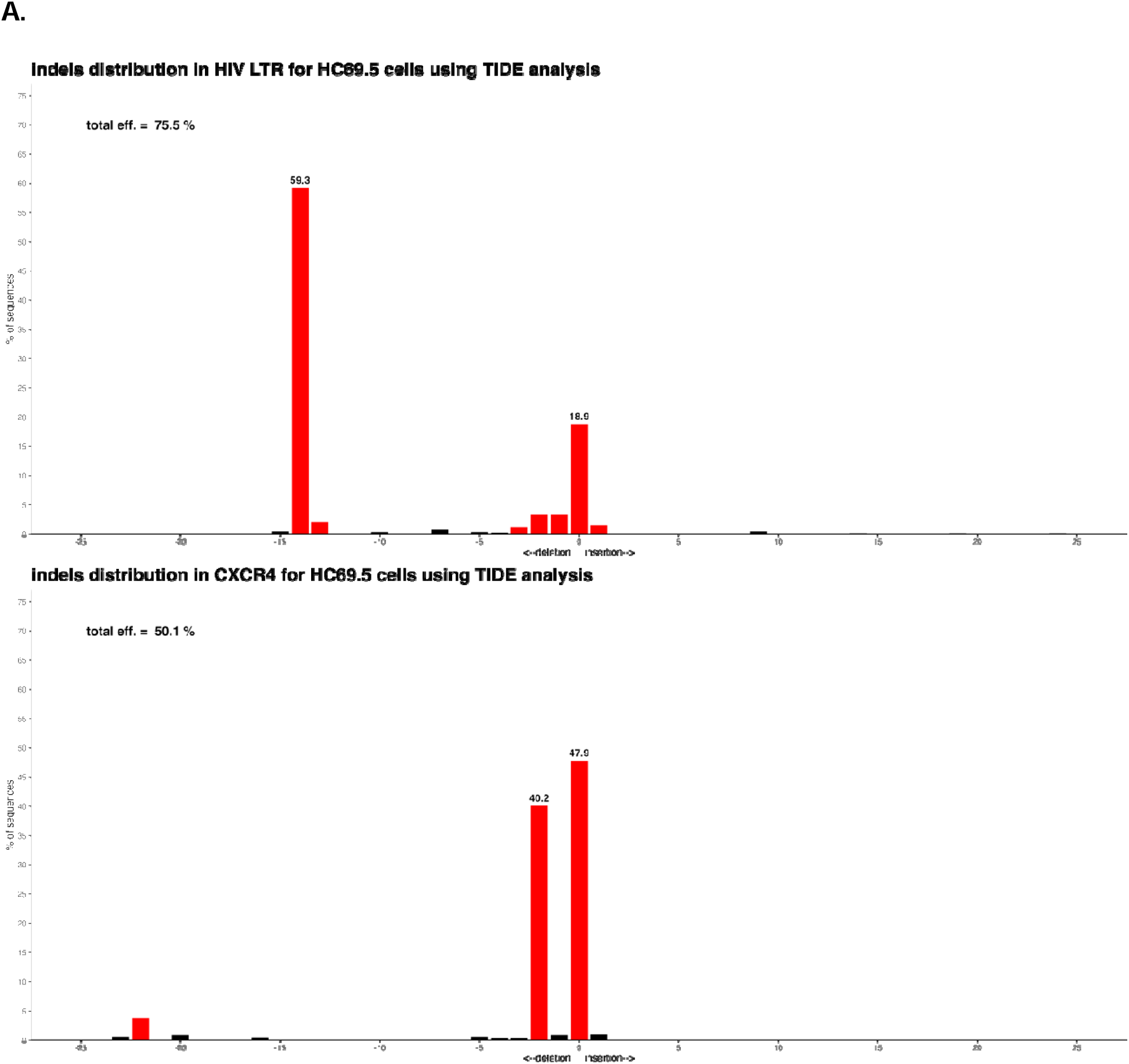
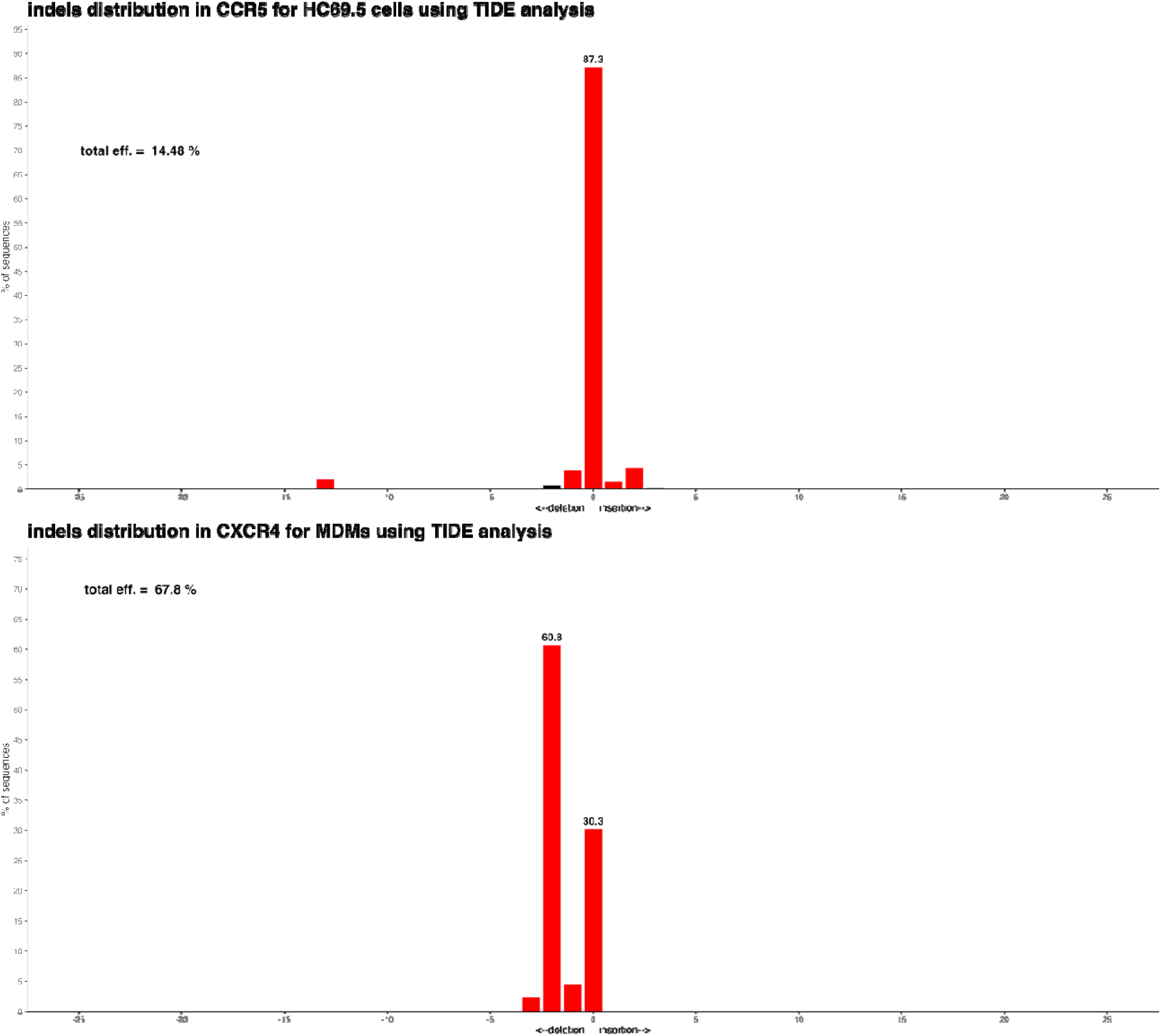
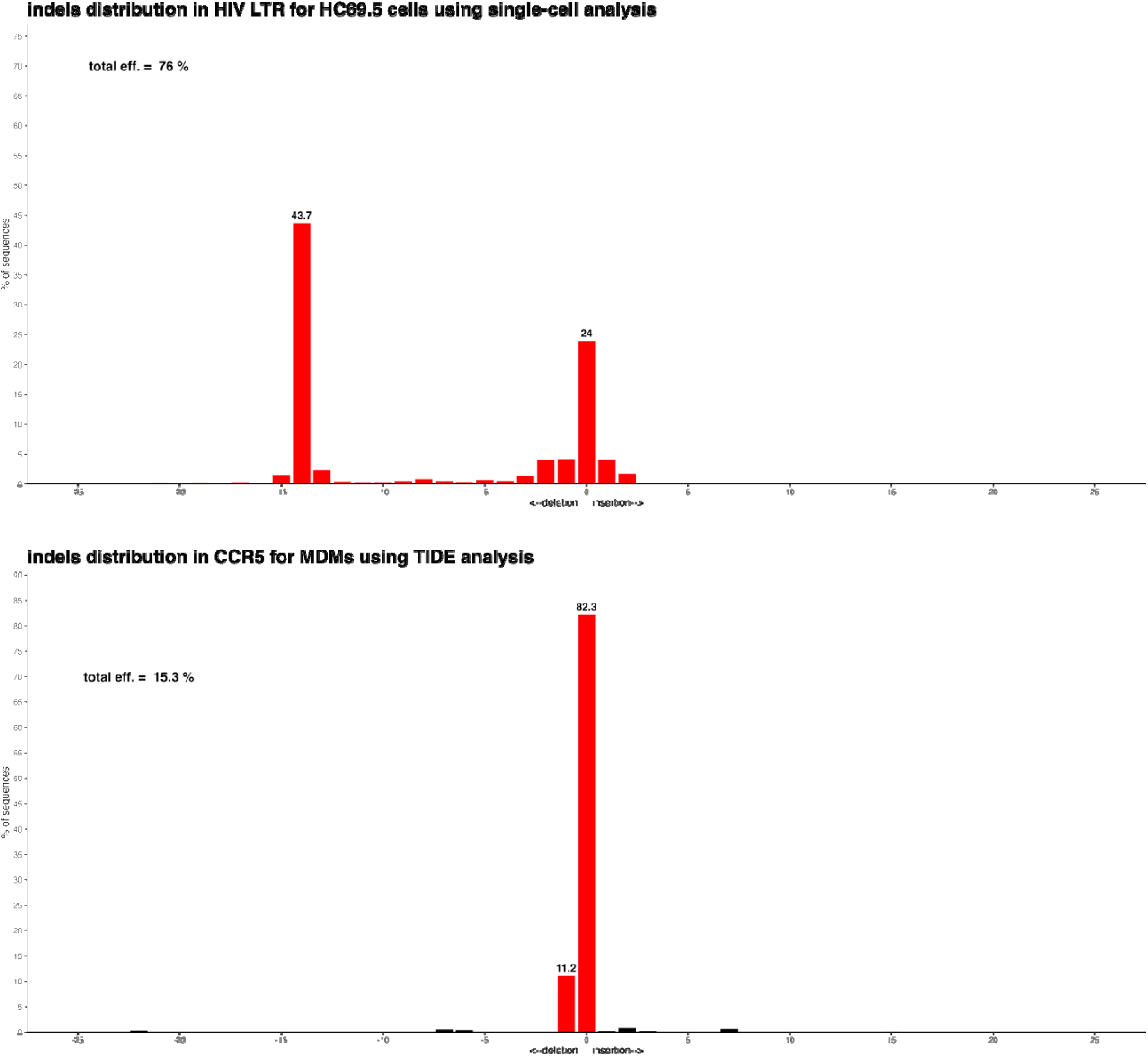
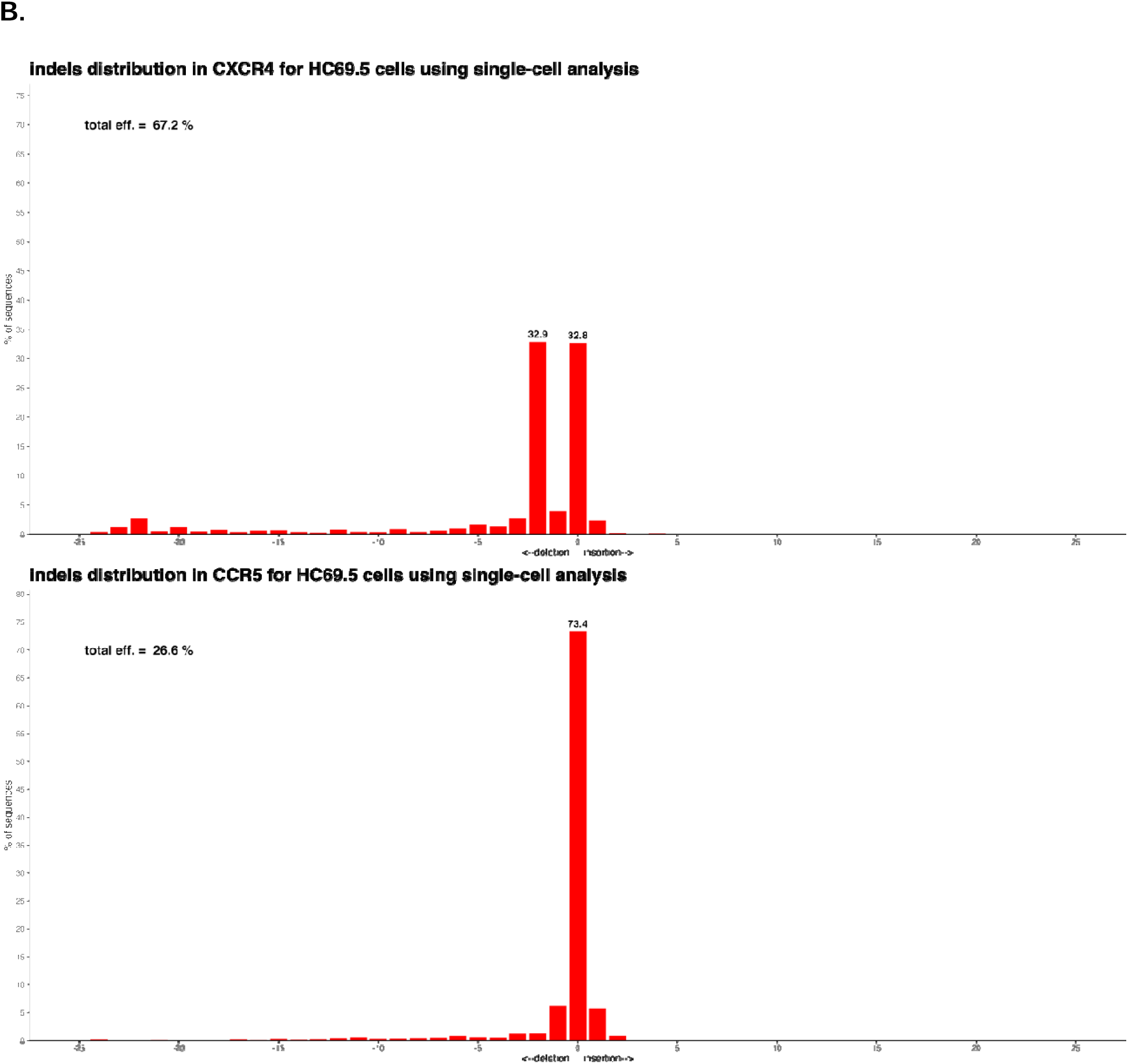
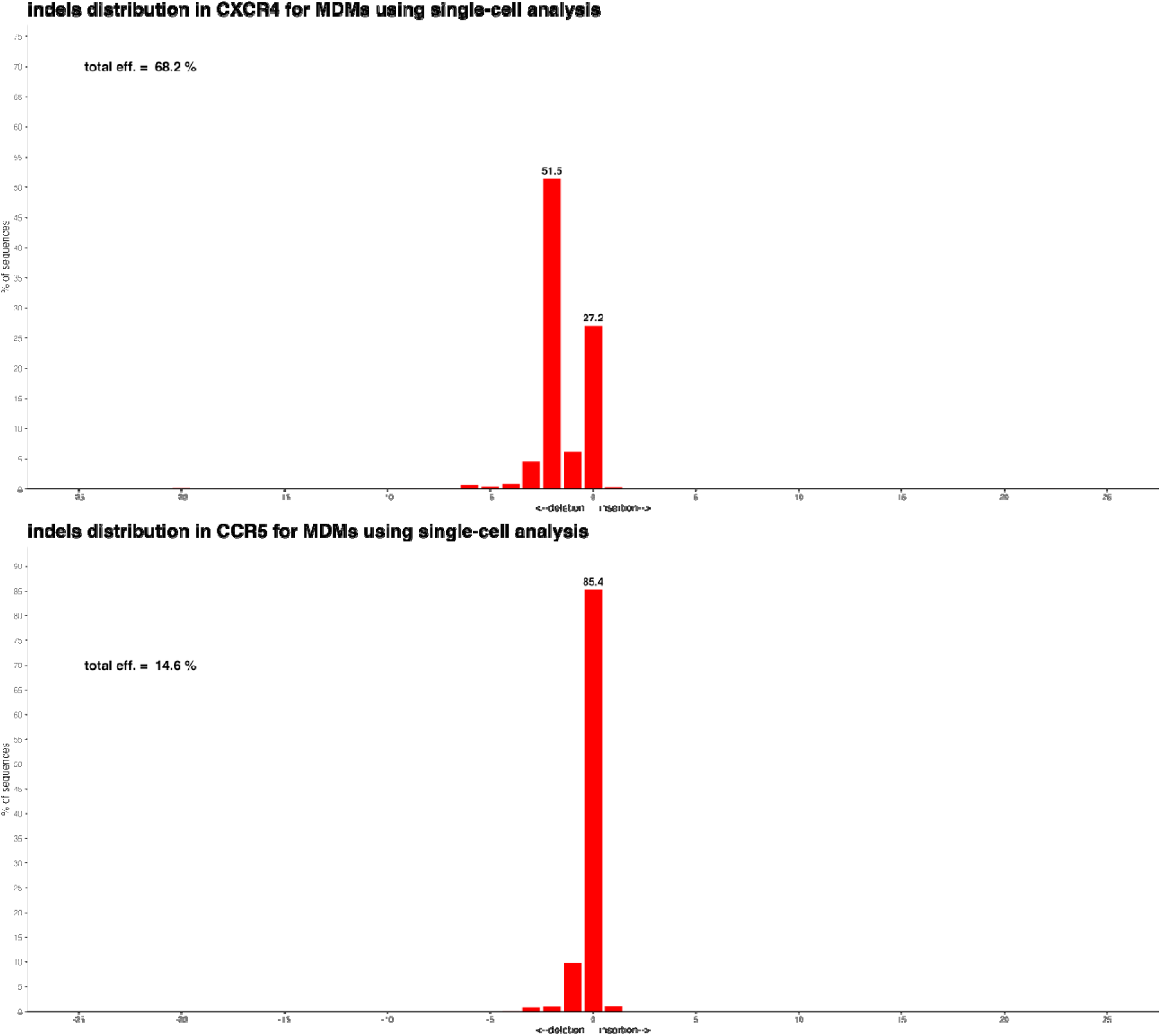
**Indel spectra in HC69.5 and MDMs infered by TIDE analysis or single cell analysis.** A) Indel spectra determined using Sanger sequencing and TIDE software. Red bars indicate indel variants occuring at statistically significant frequencies as determined by TIDE analysis (Pearson’s chi-squared p<0.001), and black bars represent indel variants that did not meet the statistical cutoff. Indels in size from -25 (25 nucleotide deletion) to +25 (25 nucleotides insertions) were plotted. Percentage contribution for each indel size is indicated above each bar (when percentages exceed 10%). B) Indel spectra characterized using our custom bioinformatic pipeline for targeted single-cell sequencing.

**Supplementary Table 1.**
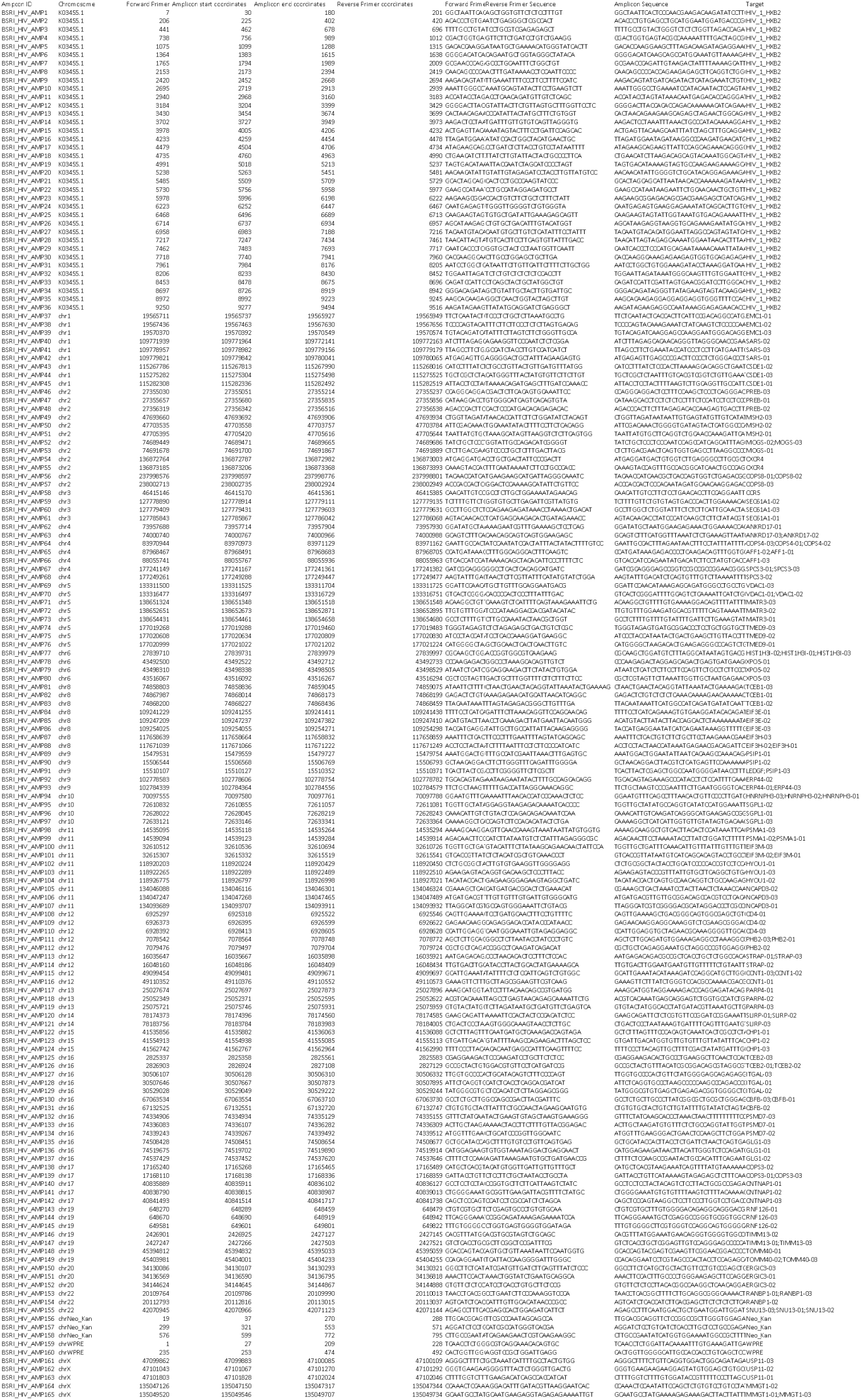
Complete list of targets in scDNA-seq panel. For each amplicon in the panel, the following information is indicated: amplicon ID, chromosome location (Genbank accession number K03455.1 pertaining to the HIV-1 HXB2 reference genome is listed for HIV amplicons), primer coordinates, forward and reverse primer sequences, amplicon sequence, and targeted gene ID are provided.

## References

1. HIV and AIDS. https://www.who.int/news-room/fact-sheets/detail/hiv-aids.

2. Eisinger, R. W., Dieffenbach, C. W. & Fauci, A. S. HIV Viral Load and Transmissibility of HIV Infection: Undetectable Equals Untransmittable. JAMA 321, 451–452 (2019).

3. Ho, Y.-C. et al. Replication-competent noninduced proviruses in the latent reservoir increase barrier to HIV-1 cure. Cell 155, 540–551 (2013).

4. Abana, C. Z.-Y., Lamptey, H., Bonney, E. Y. & Kyei, G. B. HIV cure strategies: which ones are appropriate for Africa? Cell Mol Life Sci 79, 400 (2022).

5. Ismail, S. D. et al. Addressing an HIV cure in LMIC. Retrovirology 18, 21 (2021).

6. Deeks, S. G. HIV: Shock and kill. Nature 487, 439–440 (2012).

7. Vansant, G., Bruggemans, A., Janssens, J. & Debyser, Z. Block-And-Lock Strategies to Cure HIV Infection. Viruses 12, 84 (2020).

8. Gonzalez-Cao, M., Martinez-Picado, J., Karachaliou, N., Rosell, R. & Meyerhans, A. Cancer immunotherapy of patients with HIV infection. Clin Transl Oncol 21, 713–720 (2019).

9. Hütter, G. et al. Long-Term Control of HIV by CCR5 Delta32/Delta32 Stem-Cell Transplantation. New England Journal of Medicine 360, 692–698 (2009).

10. Gupta, R. K. et al. HIV-1 remission following CCR5Δ32/Δ32 haematopoietic stem-cell transplantation. Nature 568, 244–248 (2019).

11. Jensen, B.-E. O. et al. In-depth virological and immunological characterization of HIV-1 cure after CCR5Δ32/Δ32 allogeneic hematopoietic stem cell transplantation. Nat Med 29, 583– 587 (2023).

12. Manjunath, N., Yi, G., Dang, Y. & Shankar, P. Newer gene editing technologies toward HIV gene therapy. Viruses 5, 2748–2766 (2013).

13. Khalili, K., Kaminski, R., Gordon, J., Cosentino, L. & Hu, W. Genome editing strategies: potential tools for eradicating HIV-1/AIDS. J Neurovirol 21, 310–321 (2015).

14. Hütter, G. et al. CCR5 Targeted Cell Therapy for HIV and Prevention of Viral Escape. Viruses 7, 4186–4203 (2015).

15. Jiang, F. & Doudna, J. A. CRISPR-Cas9 Structures and Mechanisms. Annu Rev Biophys 46, 505–529 (2017).

16. Chavez, L. R. et al. Durable Control of HIV-1 Using a *Staphylococcus aureus* Cas9-Expressing Lentivirus Co-Targeting Viral Latency and Host Susceptibility. bioRxiv 2020.08.10.243329 (2020) doi:10.1101/2020.08.10.243329.

17. Yin, C. et al. In Vivo Excision of HIV-1 Provirus by saCas9 and Multiplex Single-Guide RNAs in Animal Models. Molecular Therapy 25, 1168–1186 (2017).

18. Yin, C. et al. Functional screening of guide RNAs targeting the regulatory and structural HIV-1 viral genome for a cure of AIDS. AIDS 30, 1163–1173 (2016).

19. Kaminski, R. et al. Excision of HIV-1 DNA by gene editing: a proof-of-concept in vivo study. Gene Ther. 23, 690–695 (2016).

20. Liao, H.-K. et al. Use of the CRISPR/Cas9 system as an intracellular defense against HIV-1 infection in human cells. Nat Commun 6, 6413 (2015).

21. Kaminski, R. et al. Elimination of HIV-1 Genomes from Human T-lymphoid Cells by CRISPR/Cas9 Gene Editing. Sci Rep 6, 22555 (2016).

22. Hu, W. et al. RNA-directed gene editing specifically eradicates latent and prevents new HIV-1 infection. PNAS 111, 11461–11466 (2014).

23. Hou, P. et al. Genome editing of CXCR4 by CRISPR/cas9 confers cells resistant to HIV-1 infection. Sci Rep 5, 15577 (2015).

24. Ye, L. et al. Seamless modification of wild-type induced pluripotent stem cells to the natural CCR5Δ32 mutation confers resistance to HIV infection. Proceedings of the National Academy of Sciences 111, 9591–9596 (2014).

25. Wang, W. et al. CCR5 Gene Disruption via Lentiviral Vectors Expressing Cas9 and Single Guided RNA Renders Cells Resistant to HIV-1 Infection. PLOS ONE 9, e115987 (2014).

26. Dash, P. K. et al. Sequential LASER ART and CRISPR Treatments Eliminate HIV-1 in a Subset of Infected Humanized Mice. Nat Commun 10, 2753 (2019).

27. Bella, R. et al. Removal of HIV DNA by CRISPR from Patient Blood Engrafts in Humanized Mice. Mol Ther Nucleic Acids 12, 275–282 (2018).

28. Mancuso, P. et al. CRISPR based editing of SIV proviral DNA in ART treated non-human primates. Nat Commun 11, 6065 (2020).

29. Hiatt, J. et al. Efficient generation of isogenic primary human myeloid cells using CRISPR-Cas9 ribonucleoproteins. Cell Rep 35, 109105 (2021).

30. Yaseen, M. M., Abuharfeil, N. M., Alqudah, M. A. & Yaseen, M. M. Mechanisms and Factors That Drive Extensive Human Immunodeficiency Virus Type-1 Hypervariability: An Overview. Viral Immunol 30, 708–726 (2017).

31. Johnson, M. M., Jones, C. E. & Clark, D. N. The Effect of Treatment-Associated Mutations on HIV Replication and Transmission Cycles. Viruses 15, 107 (2022).

32. Wang, G., Zhao, N., Berkhout, B. & Das, A. T. CRISPR-Cas9 Can Inhibit HIV-1 Replication but NHEJ Repair Facilitates Virus Escape. Mol Ther 24, 522–526 (2016).

33. De Silva Feelixge, H. S., et al. Detection of treatment-resistant infectious HIV after genome-directed antiviral endonuclease therapy. Antiviral Res 126, 90–98 (2016).

34. Ueda, S., Ebina, H., Kanemura, Y., Misawa, N. & Koyanagi, Y. Anti-HIV-1 potency of the CRISPR/Cas9 system insufficient to fully inhibit viral replication. Microbiology and Immunology 60, 483–496 (2016).

35. Wang, Z. et al. CRISPR/Cas9-Derived Mutations Both Inhibit HIV-1 Replication and Accelerate Viral Escape. Cell Rep 15, 481–489 (2016).

36. Yoder, K. E. & Bundschuh, R. Host Double Strand Break Repair Generates HIV-1 Strains Resistant to CRISPR/Cas9. Sci Rep 6, 29530 (2016).

37. Sei, S. et al. Protective effect of CCR5 delta 32 heterozygosity is restricted by SDF-1 genotype in children with HIV-1 infection. AIDS 15, 1343–1352 (2001).

38. Trecarichi, E. M. et al. Partial protective effect of CCR5-Delta 32 heterozygosity in a cohort of heterosexual Italian HIV-1 exposed uninfected individuals. AIDS Res Ther 3, 22 (2006).

39. Huang, Y. et al. The role of a mutant CCR5 allele in HIV-1 transmission and disease progression. Nat Med 2, 1240–1243 (1996).

40. Bennett, E. P. et al. INDEL detection, the ‘Achilles heel’ of precise genome editing: a survey of methods for accurate profiling of gene editing induced indels. Nucleic Acids Research 48, 11958–11981 (2020).

41. Hiatt, J. et al. A functional map of HIV-host interactions in primary human T cells. Nat Commun 13, 1752 (2022).

42. Burdo, T. H. et al. Preclinical safety and biodistribution of CRISPR targeting SIV in non-human primates. Gene Ther 1–10 (2023) doi:10.1038/s41434-023-00410-4.

43. Garcia-Mesa, Y. et al. Immortalization of primary microglia: a new platform to study HIV regulation in the central nervous system. J Neurovirol 23, 47–66 (2017).

44. Yndart Arias, A., et al. Anti-inflammatory effects of CBD in human microglial cell line infected with HIV-1. Sci Rep 13, 7376 (2023).

45. Veenhuis, R. T. et al. Monocyte-derived macrophages contain persistent latent HIV reservoirs. Nat Microbiol 1–12 (2023) doi:10.1038/s41564-023-01349-3.

46. Chakrabarti, A. M. et al. Target-Specific Precision of CRISPR-Mediated Genome Editing. Molecular Cell 73, 699–713.e6 (2019).

47. Regoes, R. R. & Bonhoeffer, S. The HIV coreceptor switch: a population dynamical perspective. Trends Microbiol 13, 269–277 (2005).

48. ten Hacken, E. et al. High throughput single-cell detection of multiplex CRISPR-edited gene modifications. Genome Biology 21, 266 (2020).

49. Kruize, Z. & Kootstra, N. A. The Role of Macrophages in HIV-1 Persistence and Pathogenesis. Front. Microbiol. 10, (2019).

50. Huang, J., Zhou, Y., Li, J., Lu, A. & Liang, C. CRISPR/Cas systems: Delivery and application in gene therapy. Front Bioeng Biotechnol 10, 942325 (2022).

51. Liu, Z. et al. Genome editing of the HIV co-receptors CCR5 and CXCR4 by CRISPR-Cas9 protects CD4+ T cells from HIV-1 infection. Cell Biosci 7, 47 (2017).

52. Yu, S. et al. Simultaneous Knockout of CXCR4 and CCR5 Genes in CD4+ T Cells via CRISPR/Cas9 Confers Resistance to Both X4- and R5-Tropic Human Immunodeficiency Virus Type 1 Infection. Human Gene Therapy 29, 51–67 (2018).

53. Li, S., Holguin, L. & Burnett, J. C. CRISPR-Cas9-mediated gene disruption of HIV-1 co-receptors confers broad resistance to infection in human T cells and humanized mice. Molecular Therapy Methods & Clinical Development 24, 321–331 (2022).

54. Dudek, A. M. et al. A simultaneous knockout knockin genome editing strategy in HSPCs potently inhibits CCR5- and CXCR4-tropic HIV-1 infection. Cell Stem Cell 31, 499–518.e6 (2024).

55. Leitner, T. et al. Analysis of heterogeneous viral populations by direct DNA sequencing. Biotechniques 15, 120–127 (1993).

56. Brinkman, E. K., Chen, T., Amendola, M. & van Steensel, B. Easy quantitative assessment of genome editing by sequence trace decomposition. Nucleic Acids Research 42, e168–e168 (2014).

57. Clement, K. et al. CRISPResso2 provides accurate and rapid genome editing sequence analysis. Nat Biotechnol 37, 224–226 (2019).

